# Slow Stress-Load Accumulation Dominates BDNF-Dependent Gain in a Ten-State Computational Model of Stress Biochemistry

**DOI:** 10.64898/2026.07.15.738784

**Authors:** Intakhar Ahmad

## Abstract

**Background:** Acute stress responses are often reversible, whereas sustained stress can produce coordinated disruption across endocrine, metabolic, inflammatory, antioxidant, and neuroplastic pathways. The Tiered Stress Biochemistry Model (TSBM) is a hypothesis-generating ten-state ordinary differential-equation framework linking a stylized cortisol signal to noradrenergic drive, vitamin C, a phenomenological aldosterone/renin-angiotensin drive, magnesium, normalized BDNF-related and Nrf2-related states, inflammation, and tryptophan-kynurenine metabolism.

**Methods:** Five prespecified scenarios (normal, acute, chronic, depression-like, and low-cortisol PTSD-like) were simulated for 168 hours. Analyses included local stability, output-specific sensitivity screening, structural ablation, a 400-draw Latin-hypercube scan over independent *±* 20% parameter ranges, uncertainty distributions for threshold-crossing times, a success-conditioned parameter-trade-off screen, and a wider scan in which hypothesized BDNF-feedback gain magnitudes varied log-uniformly from 0.1 to 10 times nominal. Parameters were classified as literature-derived, literature-constrained/model-defined, calibrated, or hypothesized.

**Results:** Under the specified forcing assumptions, sustained stress produced coordinated changes across several pathways. In the depression-like scenario, removing slow stress-load accumulation increased day-7 BDNF-related activity from 47.2% to 82.6%, reduced inflammation from 14.63 to 2.15 arbitrary units, and lowered KYN/TRP from 0.173 to 0.057. Removing BDNF-dependent gain increased BDNF only to 49.1% and delayed KYN/TRP crossing by 2.5 hours. The stricter multi-output conclusion was retained in 91% of uncertainty draws. Median crossing times retained the nominal sequence, but the complete four-event order occurred in only 43% of all draws.

**Conclusions:** Within this reduced model, a shared slow stress-load process coordinates the high-exposure state, whereas BDNF-dependent feedback acts mainly as an amplifier. The model generates an experimentally testable staging hypothesis: under sustained high-exposure forcing, magnesium changes may precede later BDNF-related and KYN/TRP changes. Longitudinal studies are required to determine whether this sequence occurs biologically, whether it is reversible, and whether it has clinical relevance. The simulations are not diagnostic or treatment recommendations.

## 1 Introduction

Stress responses vary across time, context, and disorder. Major depressive disorder and post-traumatic stress disorder can show different group-level hypothalamic-pituitary-adrenal (HPA) patterns, but these distributions overlap and do not define individual diagnoses (Yehuda et al., 1996). The allostasis framework emphasizes that pathology may reflect the cumulative cost of repeated or sustained mediator exposure rather than an acute response alone (McEwen and Stellar, 199 ; McEwen, 1998).

Mathematical models of the HPA axis have clarified negative feedback, circadian entrainment, and ultradian pulsatility (Walker et al., 2010; Sriram et al., 2012; Andersen et al., 201). TSBM does not replace those mechanistic HPA models. It treats cortisol as an empirical forcing signal and asks how a deliberately reduced downstream network behaves under specified inputs.

The selected couplings have varying levels of biological support. The locus coeruleus noradrenergic system is a principal stress-responsive hub whose output is gated by tyrosine hydroxylase, the rate-limiting step in catecholamine synthesis, and whose dysregulation has been associated with neuroin-flammatory and demyelinating processes (Nagatsu et al., 1964; Valentino and Van Bockstaele, 2008; Benarroch, 2009; Feinstein et al., 2016; Polak et al., 2011). Dopamine-beta-hydroxylase requires ascorbate; ascorbate additionally serves as a cofactor in the adrenal cortex and medulla, is consumed during psychological stress responses, and turns over on a timescale that constrains its recovery dynamics and its contribution to immune function (Diliberto and Allen, 1981; Bornstein et al., 200 ; Patak et al., 2004; Brody et al., 2002; Kallner et al., 1979; Levine et al., 1996; Lykkesfeldt and Tveden-Nyborg, 2019; Carr and Maggini, 2017). Stress and aldosterone-related physiology can increase magnesium loss, and insufficient magnesium status has been linked to depression- and anxiety-related phenotypes (Golf et al., 1998; Seelig, 1994; Crook et al., 2021). Sustained glucocorticoid exposure can suppress BDNF-related expression and signaling through transcriptional, receptor-level, and chromatin mechanisms (Smith et al., 1995; Duman and Monteggia, 2006; Tsankova et al., 2006; Numakawa et al., 2009; Suri and Vaidya, 201), while glucocorticoid-receptor signaling can repress Nrf2-mediated antioxidant responses that are themselves altered in depression (Alam et al., 2017; Maes et al., 2011). Finally, inflammation can increase indoleamine 2, -dioxygenase flux through the kynurenine pathway, a route repeatedly implicated in depression (Dantzer et al., 2008; Miller and Raison, 2016; Reus et al., 2015). These observations motivate term directions but do not uniquely determine functional forms or numerical coefficients.

The principal objectives are to distinguish imposed scenario behavior from structural consequences, quantify uncertainty within the selected equations, identify pathway attribution through ablation, and define measurements capable of rejecting the proposed couplings.

## 2 Methods

### 2.1 Model architecture

TSBM contains ten state variables: locus-coeruleus noradrenergic drive (NE), vitamin C (VitC), a phenomenological aldosterone/renin-angiotensin-system drive (Ald), magnesium (Mg), normalied BDNF-related and Nrf2-related states, inflammation (INF), tryptophan (Trp), kynurenine (Kyn), and a slow cortisol-elevation state (*C*_stress_). BDNF and Nrf2 are percentages of reference activity/expression, not serum concentrations. Ald and INF are arbitrary-unit proxies. VitC, Mg, Trp, and Kyn are concentration-scale model pools rather than predictions for a specific compartment.

The implemented equations are:

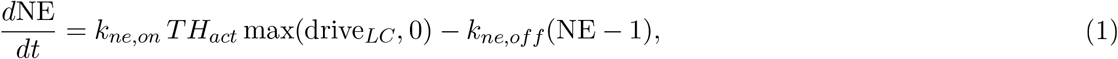

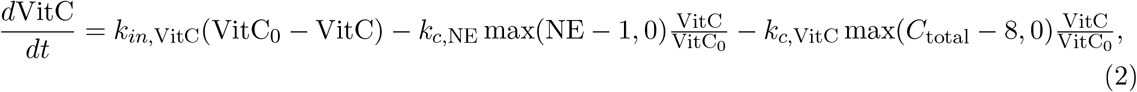

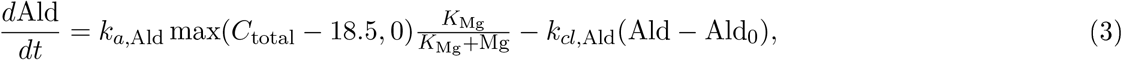

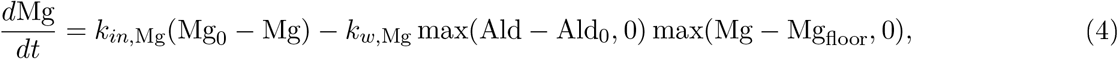

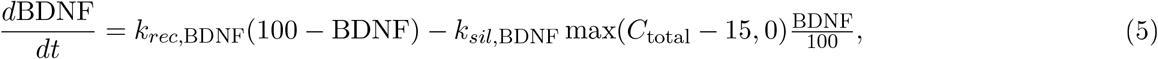

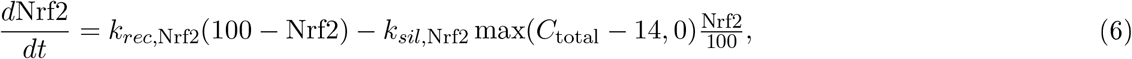

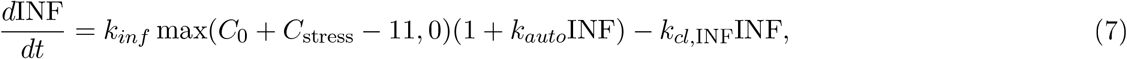

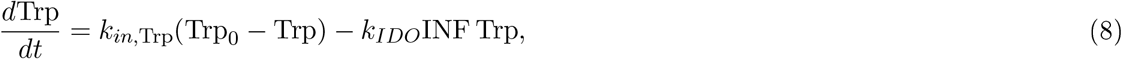

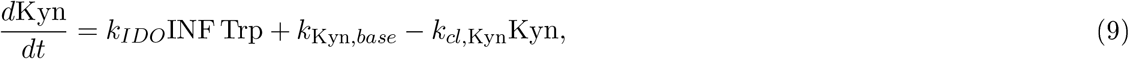

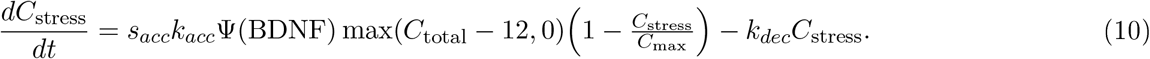

Here *s*_*acc*_ *∈ {*0, 1*}* is the scenario-specific accumulation switch and *k*_*acc*_ is the accumulation rate. This notation removes the earlier ambiguity in which “acc” was used for both concepts. *TH*_*act*_ = 1 + *k*_*thind*_*C*_stress_ and drive_*LC*_ = [*C*_acute_ + max(*C*_total_ − *C*_0_, 0)]*/*10. Full annotations and dimensional definitions are in Supplementary File S1.

### 2.2 Cortisol forcing and BDNF-dependent gain

Total cortisol is *C*_total_ = *C*_circadian_ + *C*_stress_ + *C*_acute_, with

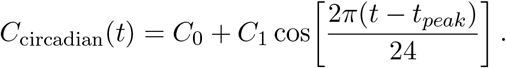

The acute pulse begins at 8 hours, has amplitude 20 *µ*g/dL, and decays with a 0.5-hour time constant. For the depression-like scenario, Ψ_dep_ = 1 + 0.60 max[0, (100 − BDNF)*/*100]; for the PTSD-like counterfactual, Ψ_PTSD_ = 1 − 0.15 max[0, (100 − BDNF)*/*100]. The exact *C*_0_*/C*_1_ pairs are literature-constrained/model-defined scenario inputs, not estimates fitted to individual participants.

### 2.3 Scenario definitions

### 2.4 Parameterization and analytical reference lines

Parameters were classified as literature-derived, literature-constrained/model-defined, calibrated, or hypothesied. A citation supports a direction, scale, or timescale and does not imply that the exact coefficient was directly measured for the modeled compartment. Analytical reference lines were Mg 0.65 mmol/L, VitC 2 *µ*M, BDNF-related state 70, and same-unit KYN/TRP 0.08. They are model references, not universal clinical cutoffs. The complete parameter table, including *k*_*in*,Trp_ and all activation thresholds, is provided in Supplementary File S2.

The display preservation index in Figure 2 was used only for coloring. Each component was clipped to [0, 1]: 1 −|NE − 1|, VitC*/*60, Mg*/*0.85, BDNF*/*100, Nrf2*/*100, 1 − INF*/*15, and 1 − (KYN*/*TRP − 0.035)*/*0.14. Raw simulated values are printed in each cell and all inferential analyses use raw outputs.

**Figure 1.**
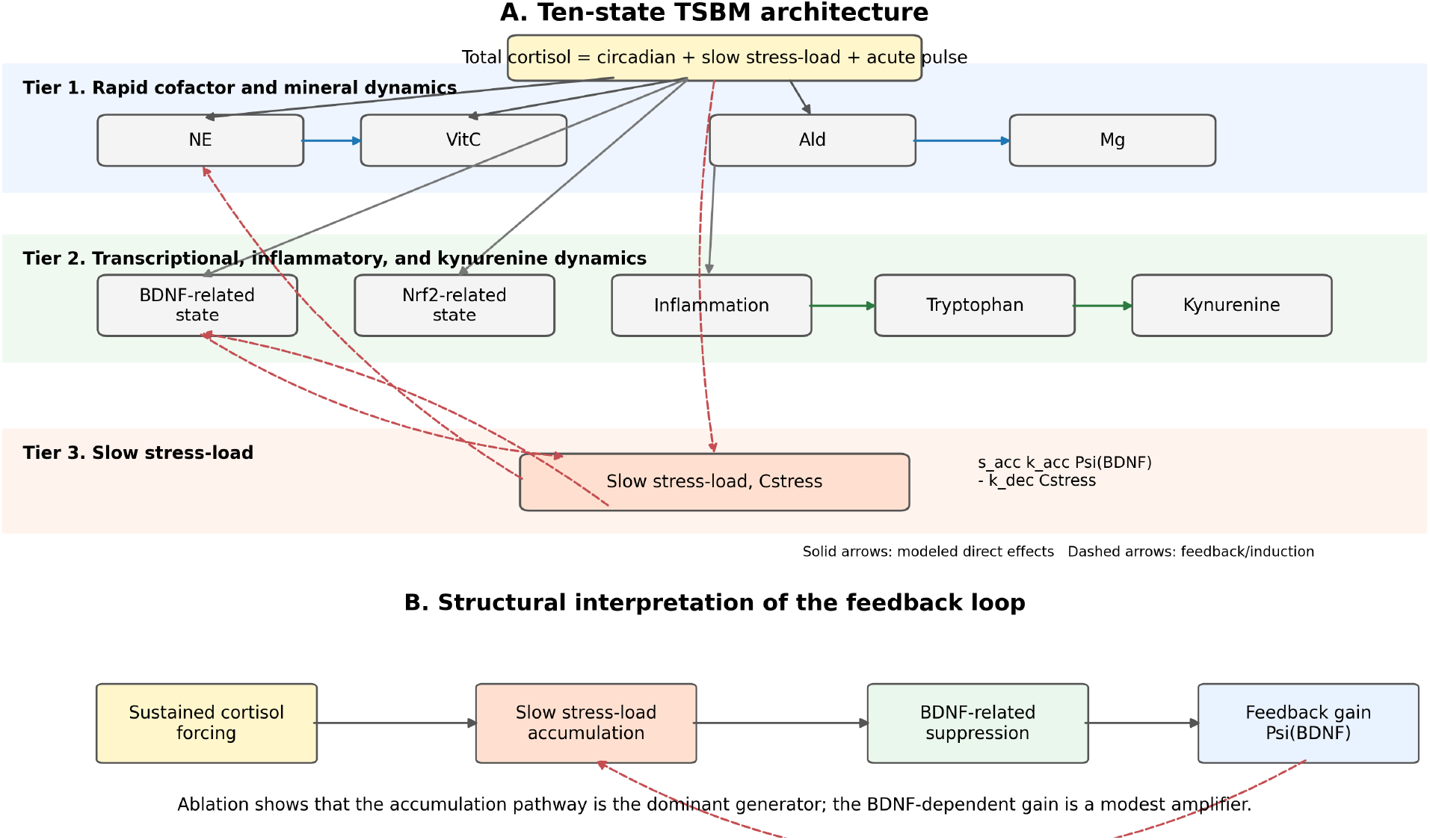
TSBM architecture and feedback interpretation. Tier 1 is labeled rapid cofactor and mineral dynamics rather than “cofactor collapse” because the acute scenario does not cross the prespecified deficiency boundaries. Structural ablation distinguishes the slow accumulation pathway from the BDNF-dependent gain.

**Table 1.** Prespecified scenario inputs. All downstream states begin at reference values except for the listed initial slow-load state.

| Scenario | $C_0$ | $C_1$ | $C_{stress}(0)$ | $s_{acc}$ | Acute pulse | $\Psi$ |
| --- | --- | --- | --- | --- | --- | --- |
| Normal | 10.0 | 7.5 | 0 | 0 | No | None |
| Acute | 10.0 | 7.5 | 0 | 0 | 20 $\mu\text{g}/\text{dL}$ | None |
| Chronic | 10.0 | 7.5 | 6 | 1 | No | None |
| Depression-like | 13.0 | 8.0 | 3 | 1 | No | Positive gain |
| Low-cortisol PTSD-like | 7.5 | 5.5 | 0 | 0 | No | Negative gain, inactive by default |

### 2.5 Numerical methods, sensitivity, uncertainty, and ablation

Main simulations used `scipy.integrate.solve_ivp` with RK45 (relative tolerance 10^−6^, absolute tolerance 10^−9^, maximum step 0.25 h). Repeated scans used a documented fixed-step RK4 implementation with 0.5-hour steps. Local sensitivity varied fourteen parameters one at a time by *±*20%. Multi-output screening evaluated day-7 VitC, Mg, BDNF, Nrf2, INF, and KYN/TRP. The normalized span was |*Y*_+20%_ − *Y*_−20%_|*/*(0.4|*Y*_nominal_|).

Global uncertainty used 400 Latin-hypercube draws over independent *±*20% ranges with seed 20240101. Threshold crossings were linearly interpolated between RK4 steps. Percentiles were calculated among crossing draws and crossing fractions were reported to retain information about right-censoring. A second 400-draw scan used seed 20240102 and additionally varied the magnitudes of both feedback gains log-uniformly from 0.1 to 10 times nominal. Spearman correlations among parameters in successful multi-output draws were interpreted as a success-conditioned trade-off screen, not identifiability. Structural ablation set major couplings to zero one at a time.

## 3 Results

### 3.1 Scenario behavior

At day 7, the normal and low-cortisol PTSD-like scenarios remained relatively preserved, whereas chronic and depression-like scenarios showed coordinated changes (Figures 2-5). This separation is conditional on the imposed baselines, accumulation switch, and initial slow-load states; it is not an emergent diagnostic classifier.

The nominal normal scenario ended at 48. *µ*M VitC and 0.85 mmol/L Mg. The low-cortisol PTSD-like comparator ended at 54.0 *µ*M and 0.85 mmol/L. The depression-like simulation ended at 2 .2 *µ*M VitC, 0.562 mmol/L Mg, 47.2 BDNF-related activity, 7.9 Nrf2-related activity, 14.6 INF units, and KYN/TRP 0.17 . Because BDNF and Nrf2 share the same recovery-plus-suppression form, their concordance is partly imposed by structure and is not independent validation.

**Figure 2.**
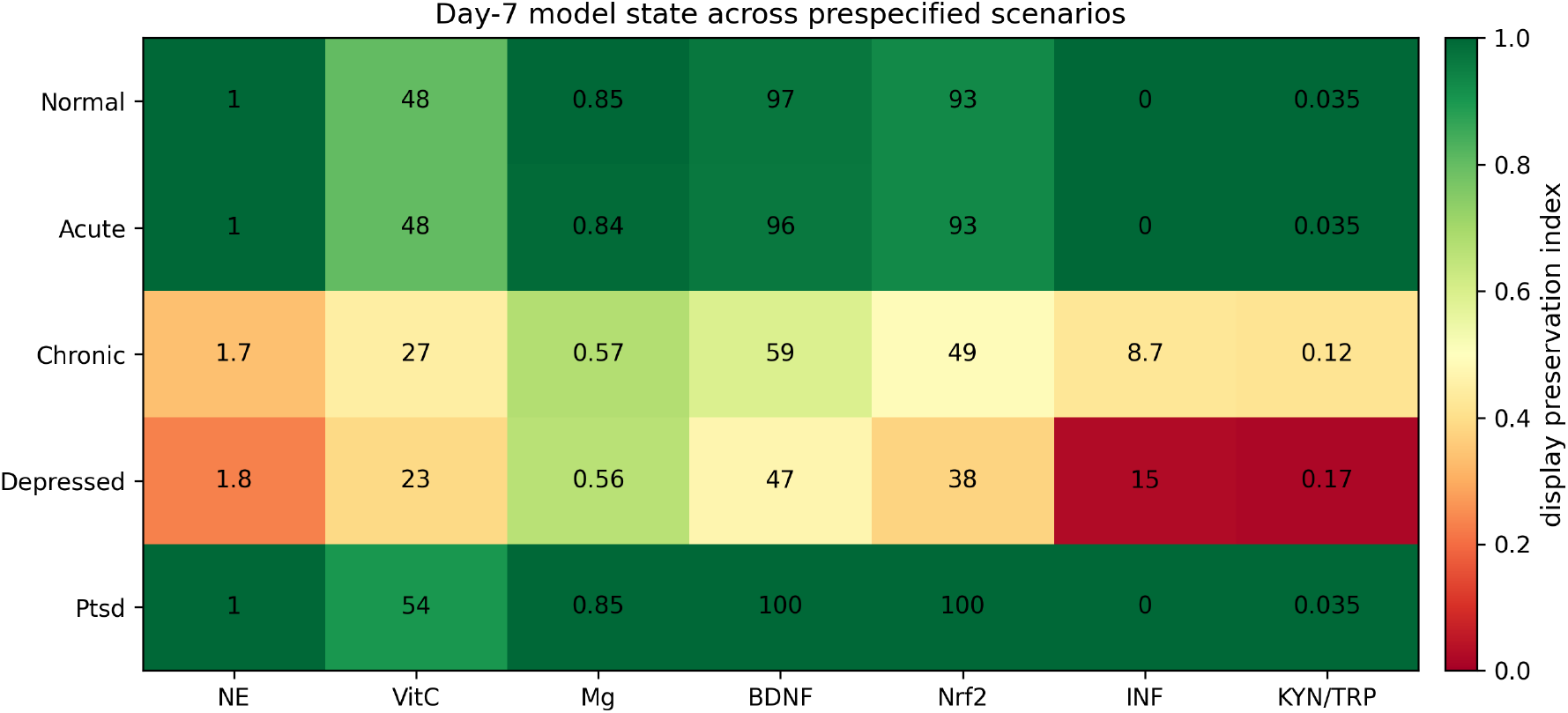
Day-7 model state across five prespecified scenarios. Color is the display-only preservation index defined in Methods; numbers are raw outputs. Scenario labels are not diagnoses.

**Figure 3.**
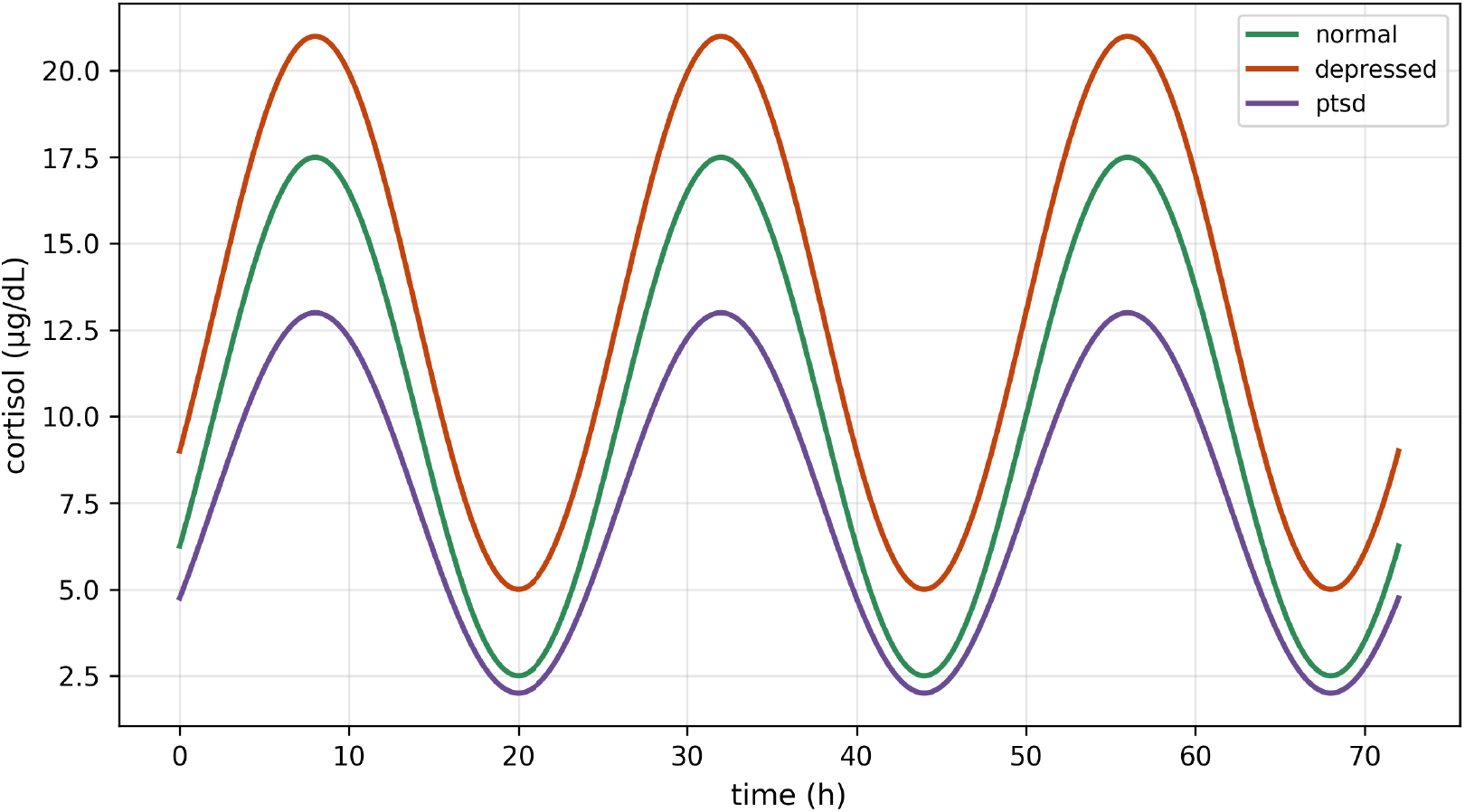
Empirical circadian cortisol backbone over 72 hours. Curves exclude slow-load and acute-pulse contributions and are scenario inputs rather than fitted patient trajectories.

**Figure 4.**
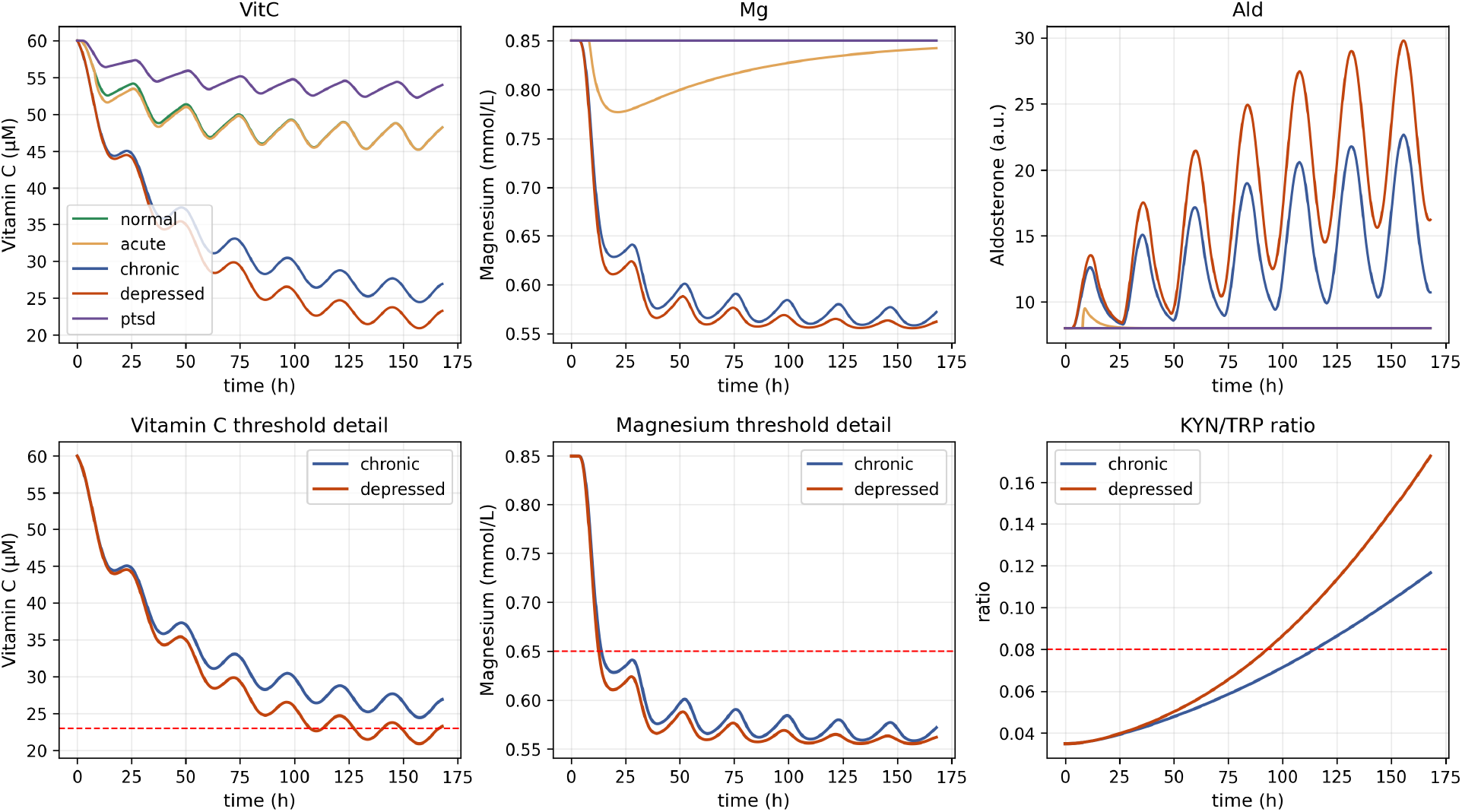
Rapid cofactor, mineral, aldosterone-proxy, and kynurenine dynamics. Dashed reference lines are analytical model boundaries rather than diagnostic cutoffs.

**Figure 5.**
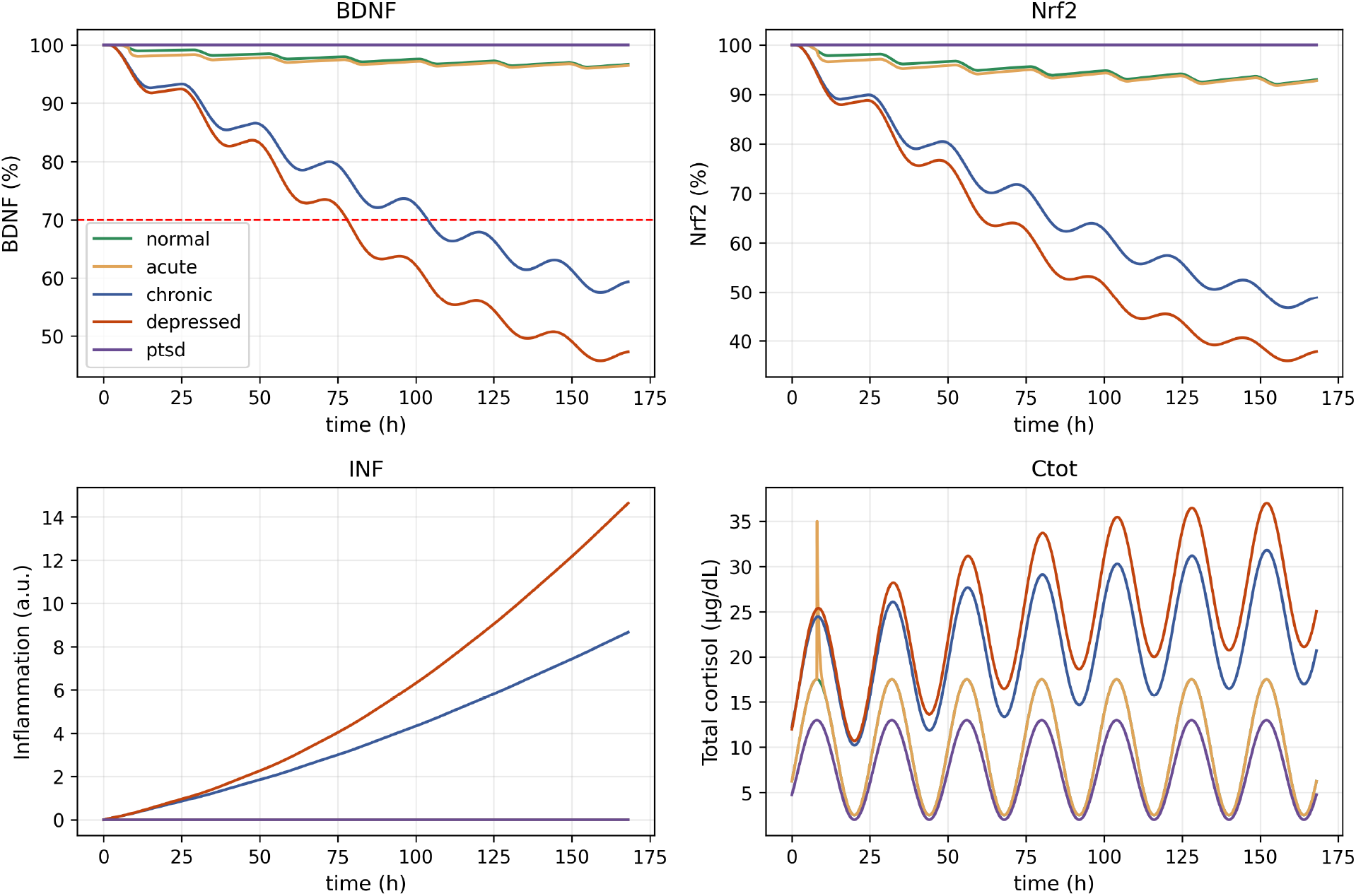
Normalized BDNF- and Nrf2-related states, inflammatory proxy, and total model cortisol. BDNF and Nrf2 are dimensionless reference-state percentages.

**Figure 6.**
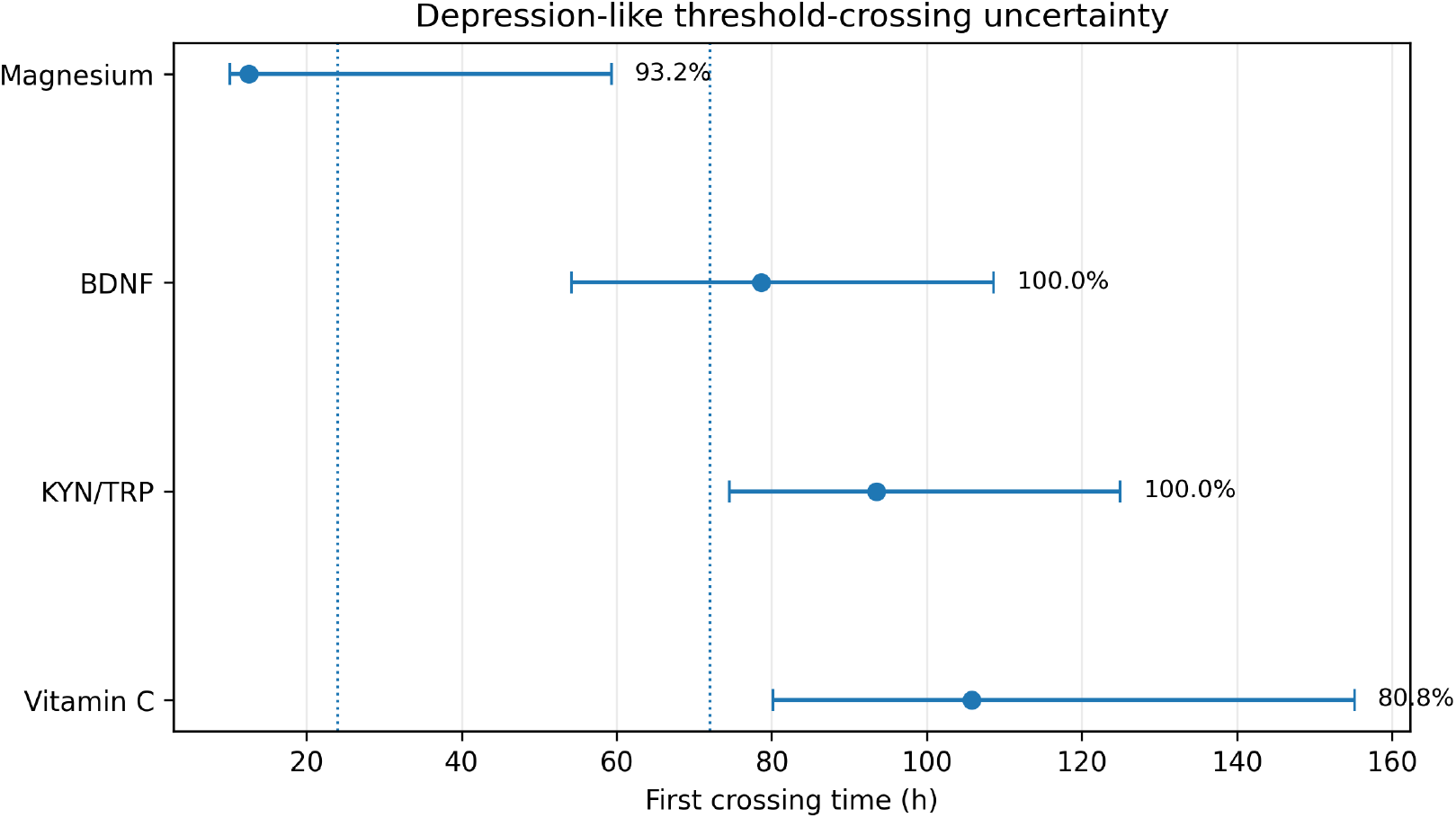
Depression-like threshold-crossing uncertainty. Points are medians and bars are 2.5th-97.5th percentiles among crossing draws; labels give crossing fractions. Non-crossers are right-censored at 168 h.

### 3.2 Threshold uncertainty and event order

The nominal depression-like trajectory crossed Mg at 12.7 h, BDNF 70 at 78.1 h, KYN/TRP 0.08 at 9 .0 h, and VitC 2 *µ*M at 107.2 h. Among crossing draws, median times (2.5th-97.5th percentiles) were 12.5 (10.1-59.) h for Mg, 78.6 (54.1-108.6) h for BDNF, 9 .5 (74.6-124.9) h for KYN/TRP, and 105.8 (80.2-155.1) h for VitC. Crossing fractions were 9 .2, 100, 100, and 80.8, respectively. All four boundaries were crossed in 0 /400 draws; the complete nominal order occurred in 172/400 draws (4 .0) and 172/ 0 complete-crossing draws (56.8).

### 3.3 Finite-horizon boundary, sensitivity, and robustness

Increasing initial slow load produced a smooth reduction in day-7 BDNF-related activity, crossing the model-defined 60 line near 4.5 *µ*g/dL (Figure 7). This is an operational finite-horion boundary, not a bifurcation or clinical threshold.

**Figure 7.**
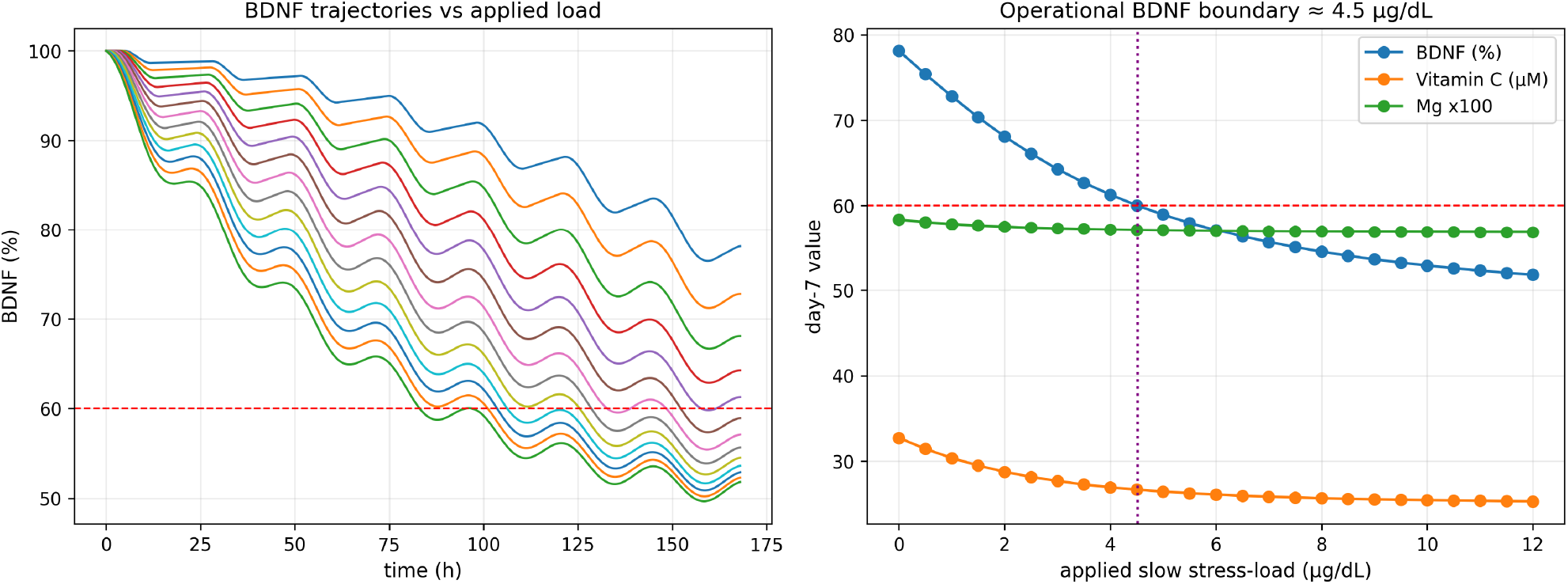
Finite-horizon operational boundary. The 60% line and approximately 4.5 *µ*g/dL crossing are analytical references.

**Figure 8.**
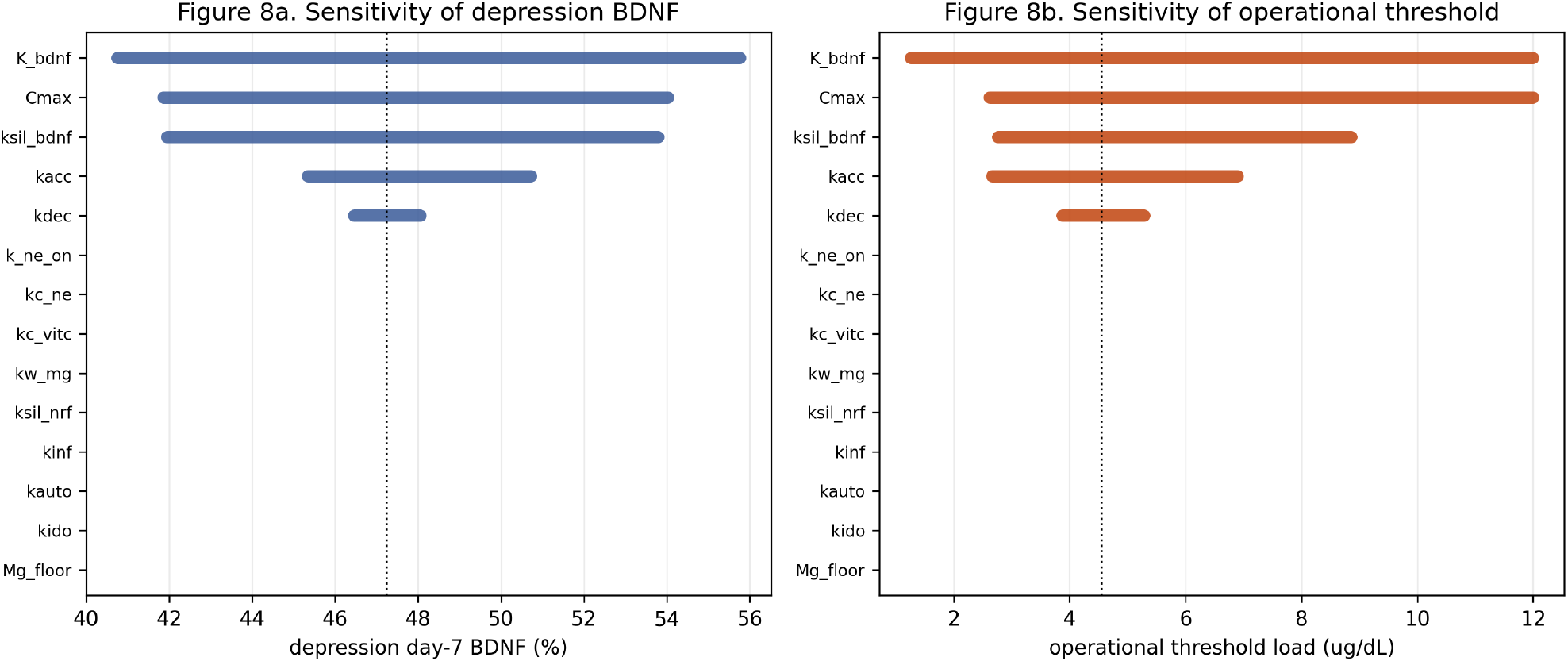
Targeted *±* 20% sensitivity of depression-like day-7 BDNF and the finite-horizon operational boundary. The apparent zero influence of some parameters is output-specific; the six-output screen is reported in Supplementary File S2.

**Figure 9.**
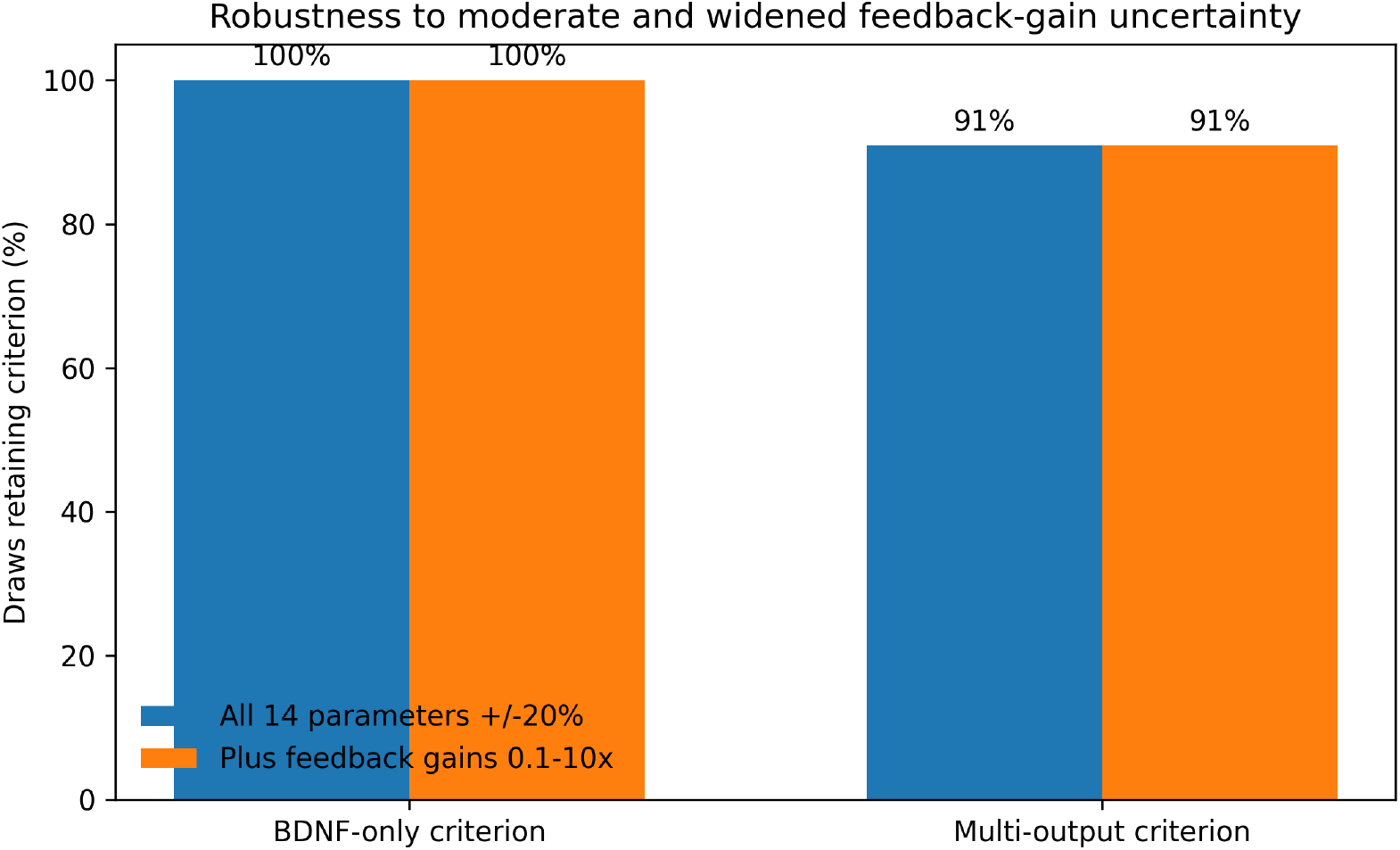
Robustness under moderate parameter uncertainty and widened feedback-gain uncertainty. The scan evaluates the selected equations and ranges, not structural uncertainty.

**Figure 10.**
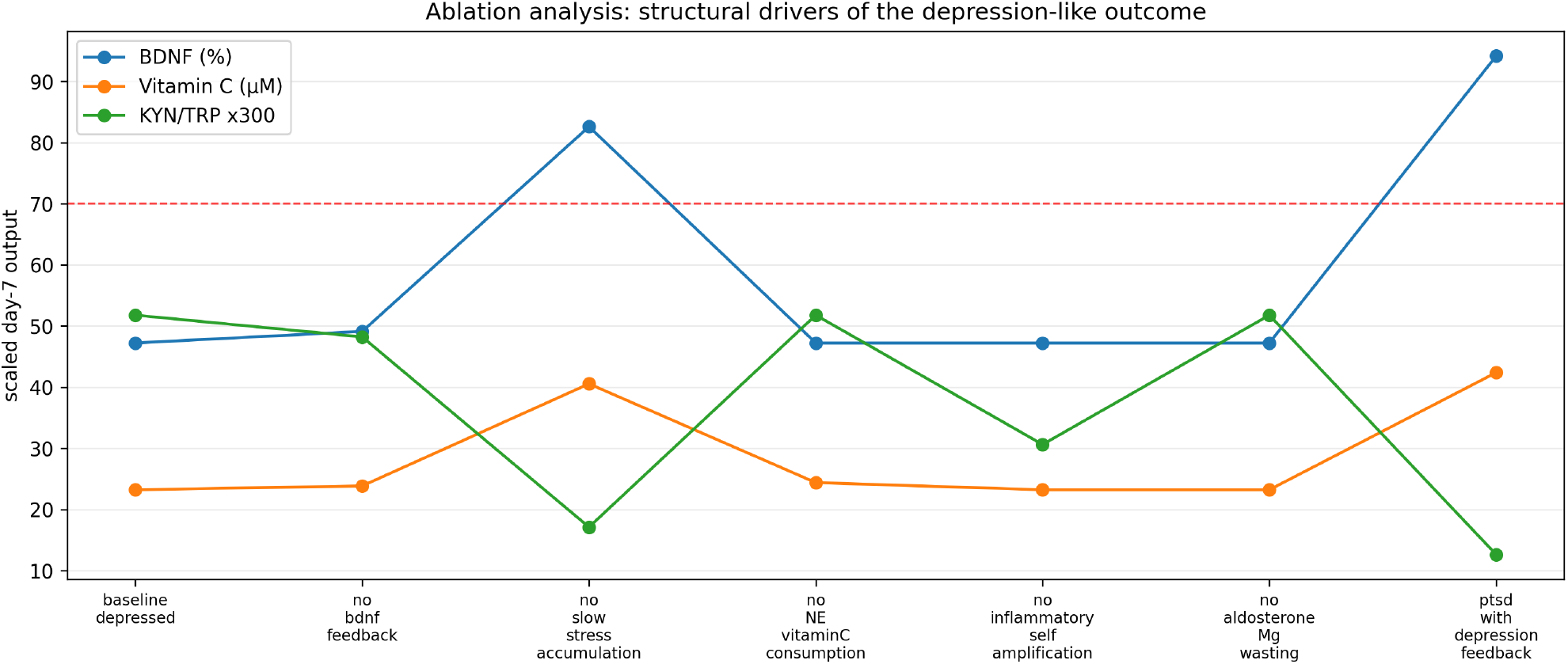
Structural-ablation analysis. Slow-load accumulation is the dominant shared coordinator, whereas the BDNF gain, inflammatory autocatalysis, and Ald-dependent Mg loss have more localized effects.

For BDNF, the largest local spans were produced by *K*_BDNF_, *C*_max_, and *k*_*sil*,BDNF_. The multi-output screen showed distinct control structure: *k*_*c*,VitC_ primarily controlled VitC; Mgfloor controlled Mg; *k*_*sil*,Nrf2_ controlled Nrf2; *k*_*inf*_ and *k*_*auto*_ controlled INF and KYN/TRP; and *C*_max_ affected all six outputs. Complete raw and normalized results are in Table S2.5.

The BDNF-only criterion was retained in 100 of moderate-uncertainty draws and the stricter multi-output criterion in 91 . These fractions were unchanged when the feedback-gain magnitudes varied over a 100-fold range. Among the 64 successful multi-output draws, the largest absolute pairwise Spearman correlation was 0.1, providing no evidence of a strong pairwise trade-off within this screen.

### 3.4 Structural ablation

Removing slow stress-load accumulation increased day-7 BDNF from 47.2 to 82.6, reduced INF from 14.6 to 2.15, and lowered KYN/TRP from 0.17 to 0.057. Removing BDNF-dependent gain increased BDNF only to 49.1 . Removing inflammatory self-amplification reduced INF to 6.80 and KYN/TRP to 0.102 without changing BDNF. Removing Ald-dependent Mg loss preserved Mg at 0.85 mmol/L. These are internal structural properties, not causal estimates from experimental data.

## 4 Discussion

TSBM is a reduced systems model for examining how a stylized endocrine input propagates through selected resource, transcriptional, inflammatory, and tryptophan-kynurenine states. The analysis separates imposed scenario behavior, calibrated consistency, uncertainty within the selected equations, and structural attribution by ablation. None constitutes out-of-sample validation.

Two conclusions are comparatively strong within the implemented structure. First, slow stress-load accumulation is substantially more influential than the hypothesied BDNF-dependent gain. Second, the low-cortisol PTSD-like scenario is preserved because exposure and accumulation are set low, not because a PTSD-specific feedback mechanism emerges. By contrast, the complete threshold sequence is not robust enough to support a treatment window: median times retain the order, but only 4 of all draws reproduce the full sequence.

The model generates falsifiable hypotheses: preventing slow-load accumulation should rescue several downstream states more strongly than removing BDNF gain alone; inflammatory self-amplification should alter KYN/TRP more than BDNF; disabling Ald-dependent loss should prevent the modeled Mg decline; and under sustained high-exposure forcing Mg frequently changes before BDNF and KYN/TRP. These hypotheses do not justify supplementation, diagnostic testing, cardiovascular risk prediction, or psychiatric treatment decisions.

Acute human stress may transiently increase peripheral BDNF, whereas Equation 5 represents only a sustained cortisol-associated suppressive component (Linz et al., 2019; Herhaus et al., 2024). Peripheral BDNF measurements in PTSD are themselves heterogeneous across studies (Mojtabavi et al., 2020), which reinforces that the normalized BDNF-related state used here should not be equated with any single circulating measurement. A future model should add an acute production term only when longitudinal data can identify it separately from recovery and suppression. Future extensions may also add CRH/ACTH, renal Mg handling, antioxidant-network interactions, or neural-tissue outcomes such as oligodendrocyte-lineage and myelin measures (Antontseva et al., 2020; Kokkosis et al., 2022), which can be quantified with standardied axon-myelin morphometry (Ahmad et al., 2025). Sustained cortisol elevation additionally produces metabolic effects such as insulin resistance (Rizza et al., 1982); a substrate tier coupling glucose and insulin was deliberately excluded here (Supplementary File S2) but is a plausible candidate for a later extension once matched longitudinal data are available. Such additions should be evaluated using held-out data or information criteria rather than improved fit to the existing calibration targets.

A prospective validation study should measure cortisol and, where feasible, ACTH, renin, and aldosterone together with Mg, VitC, BDNF, Nrf2- or oxidative-response markers, inflammatory markers, Trp, and Kyn at baseline and approximately 6, 12, 24, 48, 72, 120, and 168 h after a standardized stress paradigm or during a defined sustained exposure. Parameters should be estimated in a training cohort with a prespecified observation model and identifiability analysis, then tested in a held-out cohort. Reproducing calibration targets is not validation.

### Limitations

Cortisol is empirical rather than generated by a mechanistic HPA subsystem. Several couplings are phenomenological. BDNF and Nrf2 are normalized proxies with similar equation forms. The PTSD-like feedback is inactive by default. Mg_floor_ materially controls Mg. The uncertainty design assumes independent parameter ranges and does not represent structural uncertainty, correlated priors, measurement error, medication effects, sex or age effects, or stochastic biological variability. Threshold percentiles are conditional on crossing. The model has not been fitted to an independent longitudinal multianalyte dataset.

## 5 Conclusions

Within the implemented equations, slow stress-load accumulation is the dominant simulated coordinator and BDNF-dependent gain is a modest amplifier. The stricter multi-output conclusion persists in 91 of tested draws, whereas the complete threshold sequence is substantially less robust. Independent longitudinal calibration, formal identifiability analysis, and held-out validation are required before diagnostic, prognostic, reversibility, or treatment-timing claims are considered.

## Abbreviations

BDNF: brain-derived neurotrophic factor
DBH: dopamine-beta-hydroxylase
HPA: hypothalamic-pituitary-adrenal
IDO: indoleamine 2, -dioxygenase
KYN: kynurenine
LC: locus coeruleus
NE: noradrenergic drive
Nrf2: nuclear factor erythroid 2-related factor 2
ODE: ordinary differential equation
PTSD: post-traumatic stress disorder
TH: tyrosine hydroxylase
TRP: tryptophan.

## Author contributions

I.A. conceived the study, developed and implemented the model, performed the analyses, generated the figures, and wrote the manuscript.

## Ethics approval

Not applicable. This computational study used no human participants, animals, or identifiable data.

## Data and code availability

The core model and analysis scripts are publicly available at https://github.com/Intakhar-Ahmad/tiered-stress-biochemistry-model. The complete reproducibility package supplied with this revised version includes the extended threshold-uncertainty, event-order, widened-Ψ, and success-conditioned parameter-trade-off analyses together with machine-readable outputs.

## Reproducibility statement

Main simulations use RK45 with relative tolerance 10^−6^, absolute tolerance 10^−9^, and maximum step 0.25 h. Repeated scans use fixed-step RK4 with 0.5-hour steps. Seeds are 20240101 and 20240102. Threshold crossings are linearly interpolated between time steps. The supplied run manifest records Python .1 .5, NumPy 2. .5, and SciPy 1.17.0 for the verified execution environment.

## Supplementary materials

Supplementary File S1: annotated equations, biological interpretation, derived quantities, and complete symbol definitions.

Supplementary File S2: complete parameter and scenario tables, numerical methods, thresholds, local and multi-output sensitivity, stability, uncertainty, robustness, trade-off, and excluded-variable analyses.

Reproducibility package: source scripts, fixed-seed extended analyses, run manifests, and CSV/JSON outputs.

