## Supplementary File S1 for "Slow Stress-Load Accumulation Dominates BDNF-Dependent Gain in a Ten-State Computational Model of Stress Biochemistry"

Slow Stress-Load Accumulation Dominates BDNF-Dependent Gain in a Ten-State Computational Model of Stress Biochemistry

Intakhar Ahmad

#### Equations, dimensional definitions, and biological interpretation

### S1.1 Model scope and notation

The state vector is $[NE, VitC, Ald, Mg, BDNF, Nrf2, INF, Trp, Kyn, C_{stress}]$. $C_{stress}$ is a one-dimensional slow cortisol-elevation proxy, not a multidimensional clinical allostatic-load score. Ald is a phenomenological stress/renin-angiotensin-aldosterone drive in arbitrary units. BDNF and Nrf2 are normalized activity/expression states (percent of reference), not serum concentrations. INF is a dimensionless arbitrary-unit inflammatory proxy. VitC, Mg, Trp, and Kyn use concentration-scale units for interpretability but remain model pools.

At each time point,

$$C_{total}(t) = C_{circadian}(t) + C_{stress}(t) + C_{acute}(t) + C_{extra},$$

where $C_{extra} = 0$ in all main scenarios. The operator $\max(x, 0)$ activates a term only when its argument is positive.

### S1.2 Tier 1: rapid cofactor and mineral dynamics

The tier is called "rapid cofactor and mineral dynamics" rather than "acute cofactor collapse" because the acute scenario does not cross the prespecified deficiency boundaries.

#### Equation 1. Locus-coeruleus noradrenergic drive

$$\frac{dNE}{dt} = \underbrace{k_{ne,on} TH_{act} \max(drive_{LC}, 0)}_{\text{stress-driven TH-gated synthesis}} - \underbrace{k_{ne,off}(NE - 1)}_{\text{return toward baseline}}.$$

Derived terms are

$$TH_{act} = 1 + k_{thind} C_{stress}, \qquad drive_{LC} = \frac{C_{acute} + \max(C_{total} - C_0, 0)}{10}.$$

**Interpretation.** The equation represents fast locus-coeruleus noradrenergic output. Tyrosine hydroxylase activity increases with slow stress load, while $k_{ne,off}$ returns NE toward the reference value 1. The state is a relative drive, not a plasma norepinephrine concentration.

**Symbols.** $k_{ne,on}$, synthesis gain; $k_{ne,off}$, clearance rate; $k_{thind}$, induction per unit slow load; $C_0$, scenario baseline cortisol.

#### Equation 2. Vitamin C dynamics

$$\frac{dVitC}{dt} = \underbrace{k_{in,VitC}(VitC_0 - VitC)}_{\text{recovery/intake}} - \underbrace{k_{c,NE} \max(NE - 1, 0) \frac{VitC}{VitC_0}}_{\text{noradrenergic/DBH-associated sink}}$$
$$- \underbrace{k_{c,VitC} \max(C_{total} - C_{b,VitC}, 0) \frac{VitC}{VitC_0}}_{\text{cortisol/oxidative sink}}, \qquad C_{b,VitC} = 8\ \mu\text{g/dL}.$$

**Interpretation.** The first term restores the model pool toward 60 $\mu$M. The second represents rapid ascorbate use associated with the dopamine-beta-hydroxylase/noradrenergic arm. The third is a slower stress-associated sink. These terms are bookkeeping approximations rather than tissue-specific consumption estimates.

**Dimensional note.** Because the two sink multipliers are dimensionless except for cortisol excess in the third term, $k_{c,NE}$ has units $\mu$M h$^{-1}$ and $k_{c,VitC}$ has units $\mu$M h$^{-1}$ per $\mu$g/dL.

#### Equation 3. Phenomenological aldosterone/RAAS drive

$$\frac{dAld}{dt} = \underbrace{k_{a,Ald}\max(C_{total} - 18.5, 0)\left(\frac{K_{Mg}}{K_{Mg} + Mg}\right)}_{\text{stress/RAAS activation amplified at low Mg}} - \underbrace{k_{cl,Ald}(Ald - Ald_0)}_{\text{return toward baseline}}.$$

**Interpretation.** Ald is a reduced arbitrary-unit stress/RAAS drive. The equation does not assert that circulating cortisol alone determines circulating aldosterone.

#### Equation 4. Magnesium dynamics

$$\frac{dMg}{dt} = \underbrace{k_{in,Mg}(Mg_0 - Mg)}_{\text{recovery}} - \underbrace{k_{w,Mg}\max(Ald - Ald_0, 0)\max(Mg - Mg_{floor}, 0)}_{\text{bounded Ald-dependent loss}}, \quad Mg_{floor} = 0.55 \text{ mmol/L}.$$

**Interpretation.** The bounded loss term prevents a negative pool. $Mg_{floor}$ is the dominant sensitivity control for the Mg output; therefore crossing times are strongly conditioned on this mathematical constraint and are not independently derived renal-physiology predictions.

### S1.3 Tier 2: transcriptional, inflammatory, and kynurenine dynamics

#### Equation 5. BDNF-related state

$$\frac{dBDNF}{dt} = \underbrace{k_{rec,BDNF}(100 - BDNF)}_{\text{recovery}} - \underbrace{k_{sil,BDNF}\max(C_{total} - 15, 0)\frac{BDNF}{100}}_{\text{sustained cortisol-associated suppression}}.$$

**Interpretation.** This normalized state summarizes a sustained neurotrophic activity/expression component. It does not represent serum BDNF and does not include an acute compensatory production term.

#### Equation 6. Nrf2-related state

$$\frac{dNrf2}{dt} = \underbrace{k_{rec,Nrf2}(100 - Nrf2)}_{\text{recovery}} - \underbrace{k_{sil,Nrf2}\max(C_{total} - 14, 0)\frac{Nrf2}{100}}_{\text{stress-associated suppression}}.$$

**Interpretation.** This is a reduced proxy for glucocorticoid/oxidative regulation. Equations 5 and 6 share the same recovery-plus-suppression form, so their concordant decline is partly imposed by structure and is not independent validation.

#### Equation 7. Inflammatory proxy

$$\frac{dINF}{dt} = \underbrace{k_{inf}\max(C_0 + C_{stress} - 11, 0)(1 + k_{auto}INF)}_{\text{sustained-stress activation and self-amplification}} - \underbrace{k_{cl,INF}INF}_{\text{clearance}}.$$

**Interpretation.** Only the sustained component $C_0 + C_{stress}$ drives this equation; circadian and acute pulses are deliberately excluded. The term is a phenomenological representation of chronic stress/glucocorticoid-resistance-associated inflammatory drive, not a claim that every cortisol excursion is pro-inflammatory.

#### Equation 8. Tryptophan dynamics

$$\frac{dTrp}{dt} = \underbrace{k_{in,Trp}(Trp_0 - Trp)}_{\text{recovery}} - \underbrace{k_{IDO} INF\, Trp}_{\text{inflammation-associated IDO diversion}}.$$

**Interpretation.** Tryptophan recovers toward 60 $\mu$M and is diverted toward kynurenine in proportion to the inflammatory proxy.

#### Equation 9. Kynurenine dynamics

$$\frac{dKyn}{dt} = \underbrace{k_{IDO} INF\, Trp}_{\text{inflammation-associated production}} + \underbrace{k_{Kyn,base}}_{\text{basal production}} - \underbrace{k_{cl,Kyn} Kyn}_{\text{clearance}}.$$

**Dimensional note.** Because INF is in arbitrary units and Trp is in $\mu$M, $k_{IDO}$ has units a.u.$^{-1}$ h$^{-1}$, not $\mu$M$^{-1}$ h$^{-1}$. The healthy same-unit KYN/TRP ratio is calibrated near 0.035. The value 0.08 is an assay- and unit-dependent analytical reference line rather than a universal diagnostic threshold.

### S1.4 Tier 3: accumulated slow stress-load

#### Equation 10. Slow stress-load dynamics

$$\frac{dC_{stress}}{dt} = \underbrace{s_{acc} k_{acc} \Psi(BDNF) \max(C_{total} - 12, 0)\left(1 - \frac{C_{stress}}{C_{max}}\right)}_{\text{scenario-enabled accumulation with saturation}} - \underbrace{k_{dec} C_{stress}}_{\text{decay}}.$$

**Symbol distinction.** $s_{acc}$ is the scenario-specific switch (0 or 1); $k_{acc} = 0.028$ h$^{-1}$ is the accumulation rate. They are separate quantities and must not both be denoted "acc."

The feedback functions are

$$\Psi_{dep}(BDNF) = 1 + 0.60 \max\left(0, \frac{100 - BDNF}{100}\right),$$

$$\Psi_{PTSD}(BDNF) = 1 - 0.15 \max\left(0, \frac{100 - BDNF}{100}\right).$$

**Interpretation.** The state rises only when accumulation is enabled and total cortisol exceeds 12 $\mu$g/dL. The ceiling $C_{max}$ keeps trajectories finite. The negative PTSD-like gain is essentially inactive in the default low-exposure scenario because $s_{acc} = 0$ and BDNF remains near 100%.

### S1.5 Cortisol components and scenario inputs

$$C_{circadian}(t) = C_0 + C_1 \cos\left[\frac{2\pi(t - t_{peak})}{24}\right], \qquad t_{peak} = 8 \text{ h},$$

$$C_{acute}(t) = \begin{cases} 0, & t < 8 \text{ h}, \\ A_{acute} \exp[-(t-8)/0.5], & t \geq 8 \text{ h}, \end{cases} \qquad A_{acute} = 20\ \mu\text{g/dL}.$$

The normal, acute, chronic, depression-like, and low-cortisol PTSD-like scenario inputs are listed in Table S2.2. These are standardized forcing configurations, not diagnostic representations of patients.

### S1.6 Complete symbol summary

| Symbol | Definition |
|---|---|
| $NE$ | Noradrenergic drive relative to baseline 1. |
| $VitC$ | Vitamin C model pool, baseline 60 $\mu$M. |
| $Ald$ | Phenomenological stress/RAAS drive, baseline 8 a.u. |
| $Mg$ | Serum-comparable exchangeable Mg pool, baseline 0.85 mmol/L. |
| $BDNF, Nrf2$ | Normalized activity/expression proxies, baseline 100%. |
| $INF$ | Inflammatory-state proxy, baseline 0 a.u. |
| $Trp, Kyn$ | Tryptophan and kynurenine model pools in $\mu$M. |
| $C_{stress}$ | Slow cortisol-elevation proxy in $\mu$g/dL. |
| $C_0, C_1$ | Scenario-specific circadian mean and amplitude. |
| $s_{acc}$ | Scenario-specific 0/1 accumulation switch. |
| $k_{acc}$ | Slow-load accumulation rate. |
| $\Psi(BDNF)$ | Hypothesized BDNF-dependent gain multiplier. |

### S1.7 Interpretive boundary

Every equation is a reduced hypothesis. Literature citations support biological direction or approximate timescale but do not uniquely identify the implemented coefficients. Structural ablation tests consequences within the equations; it does not establish experimental causality. Thresholds and timing outputs are analytical model references, not clinical recommendations.
