## Supplementary File S2 for "Slow Stress-Load Accumulation Dominates BDNF-Dependent Gain in a Ten-State Computational Model of Stress Biochemistry"

### Slow Stress-Load Accumulation Dominates BDNF-Dependent Gain in a Ten-State Computational Model of Stress Biochemistry


#### Intakhar Ahmad


*Complete parameters, scenario definitions, analytical boundaries, numerical methods, sensitivity analyses, robustness analyses, and structural ablations*

#### Contents


| Section | Content |
|---|---|
| S2.1 | Parameter classification and complete parameter table |
| S2.2 | Scenario definitions |
| S2.3 | Nominal analytical-boundary crossing times |
| S2.4 | Numerical methods and reproducibility |
| S2.5 | Local sensitivity analyses |
| S2.6 | Calibration targets and contextual consistency |
| S2.7 | Stability, uncertainty, robustness, and parameter trade-offs |
| S2.8 | Structural ablation summary |
| S2.9 | Excluded candidate variables |
| S2.10 | Evidence basis for model equations |

#### S2.1 Parameter classification and complete parameter table

Parameters are classified as literature-derived, literature-constrained/model-defined, calibrated, or hypothesized. A citation supports the direction, approximate scale, or timescale of a model term; it does not imply that the exact coefficient was measured in the modeled compartment. Calibrated values reproduce prespecified model targets and are not asserted to be uniquely identifiable.

| Parameter | Value | Unit | Classification | Meaning and basis |
|---|---|---|---|---|
| C0, C1 (normal) | 10, 7.5 | µg/dL | Literature-constrained/model-defined | Stylized normal circadian mean and amplitude informed by group-level chronobiology (Yehuda et al., 1996). |
| C0, C1 (depression-like) | 13, 8 | µg/dL | Literature-constrained/model-defined | Stylized elevated-baseline profile; not an individual-level fit. |
| C0, C1 (PTSD-like) | 7.5, 5.5 | µg/dL | Literature-constrained/model-defined | Stylized low-exposure comparator; not a diagnostic biomarker. |
| t_peak | 8 | h | Literature-constrained | Morning circadian peak phase. |
| k_ne,on | 2.0 | h^-1, scaled | Calibrated/literature-constrained | Noradrenergic synthesis gain. |
| k_ne,off | 4.0 | h^-1 | Literature-constrained | Fast return of NE toward its reference value of 1. |
| k_thind | 0.020 | (µg/dL)^-1 | Calibrated/literature-constrained | Tyrosine-hydroxylase activity induction by slow stress load. |
| k_in,VitC | 0.023 | h^-1 | Literature-constrained/calibrated | Recovery toward 60 µM, informed by ascorbate turnover. |
| k_c,NE | 0.190 | µM h^-1, scaled | Calibrated | Noradrenergic/DBH-associated sink multiplying NE excess and VitC/VitC0. |
| k_c,VitC | 0.105 | µM h^-1 per µg/dL, scaled | Calibrated | Stress-associated sink above Cb,VitC. |
| Cb,VitC | 8 | µg/dL | Model-defined/calibrated | Activation threshold for the cortisol/oxidative vitamin C sink. |
| k_a,Ald | 0.55 | a.u. h^-1 per µg/dL, scaled | Calibrated/literature-constrained | Stress/RAAS proxy activation gain. |
| k_cl,Ald | 0.20 | h^-1 | Literature-constrained/calibrated | Return of the Ald proxy toward baseline. |
| th_Ald | 18.5 | µg/dL | Calibrated | Activation threshold for the Ald proxy. |
| K_Mg,Ald | 0.6 | mmol/L | Calibrated | Magnesium sensitivity constant in Ald activation. |
| k_in,Mg | 0.016 | h^-1 | Calibrated/literature-constrained | Recovery toward 0.85 mmol/L. |
| k_w,Mg | 0.040 | a.u.^-1 h^-1 | Calibrated | Bounded Ald-drive-dependent magnesium loss. |
| Mg_floor | 0.55 | mmol/L | Model-defined/calibrated | Lower bound for the exchangeable magnesium proxy; dominant magnesium sensitivity control. |
| k_rec,BDNF | 0.010 | h^-1 | Calibrated | Recovery of the normalized BDNF-related state toward 100%. |
| k_sil,BDNF | 0.100 | h^-1 per µg/dL, scaled | Calibrated/literature-constrained | Cortisol-associated BDNF-related suppression. |
| K_BDNF | 15 | µg/dL | Calibrated | BDNF-related suppression threshold. |
| k_rec,Nrf2 | 0.010 | h^-1 | Calibrated | Recovery of the normalized Nrf2-related state toward 100%. |
| k_sil,Nrf2 | 0.130 | h^-1 per µg/dL, scaled | Calibrated/literature-constrained | Stress-associated Nrf2-related suppression; direction informed by Alam et al. (2017). |
| K_Nrf2 | 14 | µg/dL | Calibrated | Nrf2-related suppression threshold. |
| k_inf | 0.006 | a.u. h^-1 per µg/dL, scaled | Calibrated/literature-constrained | Sustained-stress inflammatory activation. |
| k_auto | 0.13 | a.u.^-1 | Hypothesized/calibrated | Inflammatory self-amplification. |
| k_cl,INF | 0.012 | h^-1 | Calibrated | Inflammation clearance. |
| th_INF | 11 | µg/dL | Calibrated | Threshold applied to C0 + Cstress. |
| k_in,Trp | 0.010 | h^-1 | Calibrated | Recovery toward Trp0 = 60 µM. |
| k_IDO | 0.0007 | a.u.^-1 h^-1 | Calibrated/literature-constrained | Inflammation-dependent IDO-associated conversion; dimensional unit matches INF × Trp. |

| Parameter | Value | Unit | Classification | Meaning and basis |
|---|---|---|---|---|
| k_cl,Kyn | 0.080 | h^-1 | Calibrated | Kynurenine clearance. |
| k_Kyn,base | 0.168 | µM h^-1 | Calibrated | Basal production yielding a healthy same-unit KYN/TRP ratio near 0.035. |
| s_acc | 0 or 1 | dimensionless | Model-defined | Scenario-specific switch enabling slow stress-load accumulation. |
| k_acc | 0.028 | h^-1, scaled | Calibrated | Slow stress-load accumulation rate, distinct from s_acc. |
| k_dec | 0.0035 | h^-1 | Calibrated | Slow stress-load decay. |
| th_acc | 12 | µg/dL | Calibrated | Slow-load accumulation threshold. |
| C_max | 18 | µg/dL | Model-defined/calibrated | Saturation ceiling. |
| Psi gain (depression/PTSD) | +0.60 / -0.15 | dimensionless | Hypothesized | BDNF-dependent feedback-gain magnitudes; the PTSD gain is inactive in the default scenario. |
| Baseline states | NE=1; VitC=60; Ald=8; Mg=0.85; BDNF=Nrf2=100; INF=0; Trp=60; Kyn=2.1 | mixed | Literature-constrained/calibrated | Reference initial conditions. |
| Acute pulse | 20; onset 8; decay 0.5 | µg/dL; h; h | Model-defined | Single transient stress pulse. |

### S2.2 Scenario definitions

| Scenario | C0 | C1 | Cstress(0) | s_acc | Acute pulse | Feedback |
|---|---|---|---|---|---|---|
| Normal | 10.0 | 7.5 | 0 | 0 | None | None |
| Acute | 10.0 | 7.5 | 0 | 0 | 20 µg/dL at 8 h | None |
| Chronic | 10.0 | 7.5 | 6 | 1 | None | None |
| Depression-like | 13.0 | 8.0 | 3 | 1 | None | Psi_dep, gain +0.60 |
| Low-cortisol PTSD-like | 7.5 | 5.5 | 0 | 0 | None | Psi_PTSD, gain -0.15; inactive |

*The chronic and depression-like scenarios begin with nonzero slow load but healthy downstream biochemical states. Crossing times are therefore standardized conditional transition times, not estimates of disease onset or duration.*

### S2.3 Nominal analytical-boundary crossing times

| Analytical reference boundary | Chronic | Depression-like |
|---|---|---|
| Mg < 0.65 mmol/L | 14.0 h | 12.7 h |
| BDNF-related state < 70% | 103.8 h | 78.1 h |
| VitC < 23 µM | Not crossed | 107.2 h |
| KYN/TRP > 0.08 | 114.7 h | 93.0 h |

*Normal, acute, and low-cortisol PTSD-like scenarios crossed none of these boundaries within 168 h. These lines are analytical model references rather than universal clinical cutoffs.*

#### S2.4 Numerical methods and reproducibility

Main simulations use scipy.integrate.solve_ivp with explicit RK45, relative tolerance 10^-6, absolute tolerance 10^-9, and maximum step 0.25 h over 168 h. Repeated sensitivity and uncertainty scans use a documented fixed-step RK4 helper with 0.5-h steps. Threshold-crossing times are linearly interpolated between adjacent RK4 time points. Fixed random seeds are 20240101 for the moderate-uncertainty scan and 20240102 for the widened feedback-gain scan.

The accompanying reproducibility package contains the core model, nominal simulations, figure-generation scripts, local sensitivity analysis, global robustness analysis, structural ablation analysis, threshold-uncertainty and event-order analyses, widened-feedback draws, Spearman parameter-trade-off outputs, machine-readable tables, and a run manifest. The reported analyses are deterministic when run with the documented seeds and software settings.

#### S2.5 Local sensitivity analyses

##### S2.5.1 BDNF-related state and finite-horizon operational boundary

The baseline depression-like day-7 BDNF-related state is 47.2%, and the baseline finite-horizon operational boundary is 4.55 µg/dL. Each listed parameter was varied one at a time by ±20%.

| Parameter | BDNF at -20% | BDNF at +20% | Span (points) | Boundary at -20% | Boundary at +20% |
|---|---|---|---|---|---|
| K_BDNF | 40.7 | 55.8 | 15.0 | 1.25 | 12.00 |
| C_max | 54.0 | 41.9 | 12.2 | 12.00 | 2.62 |
| k_sil,BDNF | 53.8 | 41.9 | 11.8 | 8.86 | 2.76 |
| k_acc | 50.7 | 45.3 | 5.4 | 6.90 | 2.66 |
| k_dec | 46.5 | 48.1 | 1.6 | 3.87 | 5.28 |
| k_ne,on | 47.2 | 47.2 | 0.0 | 4.55 | 4.55 |
| k_c,NE | 47.2 | 47.2 | 0.0 | 4.55 | 4.55 |
| k_c,VitC | 47.2 | 47.2 | 0.0 | 4.55 | 4.55 |
| k_w,Mg | 47.2 | 47.2 | 0.0 | 4.55 | 4.55 |
| k_sil,Nrf2 | 47.2 | 47.2 | 0.0 | 4.55 | 4.55 |
| k_inf | 47.2 | 47.2 | 0.0 | 4.55 | 4.55 |
| k_auto | 47.2 | 47.2 | 0.0 | 4.55 | 4.55 |
| k_IDO | 47.2 | 47.2 | 0.0 | 4.55 | 4.55 |
| Mg_floor | 47.2 | 47.2 | 0.0 | 4.55 | 4.55 |

##### S2.5.2 Six-output sensitivity screen

Normalized span is |Y(+20%) - Y(-20%)| / (0.4 × |Y_nominal|). The table reports output-specific sensitivity across vitamin C, magnesium, BDNF-related activity, Nrf2-related activity, inflammation, and KYN/TRP. This is a screening metric, not an identifiability estimate.

| Parameter | VitC | Mg | BDNF | Nrf2 | INF | KYN/TRP |
|---|---|---|---|---|---|---|
| k_ne,on | .048 | .000 | .000 | .000 | .000 | .000 |
| k_c,NE | .048 | .000 | .000 | .000 | .000 | .000 |
| k_c,VitC | .547 | .000 | .000 | .000 | .000 | .000 |

| Parameter | VitC | Mg | BDNF | Nrf2 | INF | KYN/TRP |
|---|---|---|---|---|---|---|
| k_w,Mg | .000 | .018 | .000 | .000 | .000 | .000 |
| k_sil,BDNF | .015 | .001 | .627 | .026 | .061 | .051 |
| K_BDNF | .022 | .002 | .795 | .044 | .109 | .093 |
| k_sil,Nrf2 | .000 | .000 | .000 | .705 | .000 | .000 |
| k_inf | .000 | .000 | .000 | .000 | 1.922 | 1.549 |
| k_auto | .000 | .000 | .000 | .000 | .901 | .681 |
| k_IDO | .000 | .000 | .000 | .000 | .000 | .846 |
| k_acc | .145 | .011 | .285 | .297 | .741 | .640 |
| k_dec | .058 | .005 | .084 | .091 | .195 | .158 |
| C_max | .504 | .046 | .644 | .716 | 1.349 | 1.039 |
| Mg_floor | .000 | .947 | .000 | .000 | .000 | .000 |

#### S2.6 Calibration targets and contextual consistency

| Quantity | Literature target or context | Model value | Interpretation |
|---|---|---|---|
| Normal cortisol peak | ~17.5 µg/dL | 17.5 | Calibration |
| Normal cortisol nadir | ~2-3 µg/dL | 2.5 | Calibration |
| Healthy same-unit KYN/TRP | 0.03-0.05 | 0.035 | Calibration |
| KYN/TRP analytical reference line | 0.08; assay- and unit-dependent | 0.173 in depression-like run | Calibration/context |
| Magnesium analytical-boundary crossing | Within 24 h; prespecified target | 13 h | Calibration |
| Normal vitamin C context | Approximately 50-70 µM | 48.3 µM | Near-range contextual consistency |
| BDNF-related suppression | 40-60% reduction | 53% reduction | Contextual consistency |

*Rows used to constrain model parameters are not independent predictive validation. The vitamin C output is described as near-range contextual consistency rather than exact calibration because 48.3 µM is slightly below the stated 50-70 µM contextual range.*

#### S2.7 Stability, uncertainty, robustness, and parameter trade-offs

The autonomous unstressed equilibrium, obtained by freezing circadian forcing and disabling accumulation, has a maximum absolute derivative below 10^-15. All Jacobian eigenvalues have negative real parts; the largest is -0.0035. This establishes local asymptotic stability only for that autonomous fixed point, not global stability of periodically forced or high-exposure scenarios.

Under 400 independent ±20% Latin-hypercube draws, the BDNF-only criterion was retained in 100% of draws and the stricter multi-output criterion in 91%. Depression-like day-7 BDNF had a median of 48.1% (2.5th-97.5th percentile, 36.3-62.3%); PTSD-like BDNF had a median of 100.0% (99.2-100.0%); and chronic BDNF had a median of 60.0% (45.6-74.9%). These results quantify robustness within the selected equations and parameter ranges, not robustness to alternative model structures.

| Boundary | Crossing fraction | Median (h) | 2.5th percentile (h) | 97.5th percentile (h) |
|---|---|---|---|---|
| Mg < 0.65 | 93.2% | 12.5 | 10.1 | 59.3 |
| BDNF < 70% | 100% | 78.6 | 54.1 | 108.6 |
| KYN/TRP > 0.08 | 100% | 93.5 | 74.6 | 124.9 |
| VitC < 23 | 80.8% | 105.8 | 80.2 | 155.1 |

All four boundaries were crossed in 303 of 400 draws. The complete nominal order occurred in 172 of 400 draws (43.0%) and in 172 of 303 complete-crossing draws (56.8%). Thus, median times retain the nominal order, but the complete sequence is not sufficiently robust to define a treatment window.

When both feedback-gain magnitudes were additionally sampled log-uniformly from 0.1 to 10 times nominal, the BDNF-only and multi-output fractions remained 100% and 91%, respectively. Among 364 moderate-uncertainty draws satisfying the multi-output criterion, the largest absolute pairwise Spearman correlation among the fourteen varied parameters was 0.129. Because this screen conditions on success and fits no likelihood, it is described as a parameter-trade-off screen rather than practical identifiability.

#### S2.8 Structural ablation summary

| Ablation | VitC (µM) | Mg (mmol/L) | BDNF (%) | INF (a.u.) | KYN/TRP | Interpretation |
|---|---|---|---|---|---|---|
| Baseline depression-like | 23.24 | 0.562 | 47.24 | 14.63 | 0.173 | Reference |
| No BDNF-dependent gain | 23.87 | 0.563 | 49.13 | 13.38 | 0.161 | Modest global effect |
| No slow stress-load accumulation | 40.57 | 0.625 | 82.62 | 2.15 | 0.057 | Dominant shared coordinator |
| No NE-associated vitamin C sink | 24.41 | 0.562 | 47.24 | 14.63 | 0.173 | Localized vitamin C effect |
| No inflammatory self-amplification | 23.24 | 0.562 | 47.24 | 6.80 | 0.102 | Controls the INF/KYN arm |
| No Ald-dependent magnesium loss | 23.24 | 0.850 | 47.24 | 14.63 | 0.173 | Controls the magnesium arm |

#### S2.9 Excluded candidate variables

| Candidate | Reason for exclusion |
|---|---|
| Serotonin | Tryptophan and kynurenine diversion are represented; adding serotonin would require synthesis, transport, and receptor parameters not identified by the current targets. |
| CRH and ACTH | Cortisol is supplied empirically; resolving upstream pituitary dynamics would require a separate mechanistic HPA subsystem. |
| Zinc and vitamin B6 | Biologically plausible cofactors, but inclusion would add unmeasured parameters and partly duplicate resource-tier behavior. |
| Selenium/glutathione peroxidase | Overlaps the reduced Nrf2 antioxidant tier. |
| DHEA | Counter-regulatory endocrine partner whose dynamics require additional states and longer-term data. |
| Telomerase | Chronic aging endpoint outside the 168-h horizon. |
| Glucose/insulin | Requires substrate, insulin, tissue-uptake, and feedback states. |

*Excluded variables are not biologically unimportant. Future additions should be justified by a specific dataset and compared with the present model using out-of-sample predictive performance or information criteria, rather than improved fit to the same calibration targets alone.*

#### S2.10 Evidence basis for model equations

| Equation/state | Biological basis | Evidence classification | Representative references |
|---|---|---|---|
| Eq. 1: NE | Tyrosine hydroxylase is rate-limiting for catecholamine synthesis; stress inputs alter LC output. | Literature-constrained direction and timescale | Nagatsu et al., 1964; Valentino & Van Bockstaele, 2008; Benarroch, 2009 |
| Eq. 2: Vitamin C | Dopamine-beta-hydroxylase is ascorbate-dependent; vitamin C turnover informs recovery timescale. | Literature-constrained structure; calibrated coefficients | Diliberto & Allen, 1981; Kallner et al., 1979; Lykkesfeldt & Tveden-Nyborg, 2019 |
| Eq. 3: Ald proxy | Stress/RAAS physiology and magnesium interactions motivate the direction of the reduced proxy. | Phenomenological, literature-constrained | Golf et al., 1998; Seelig, 1994; Crook et al., 2021 |
| Eq. 4: Magnesium | Stress-associated mineral loss motivates Ald-dependent depletion; the floor is mathematical. | Literature-constrained direction; calibrated bounded form | Golf et al., 1998 |
| Eq. 5: BDNF | Sustained glucocorticoid exposure can suppress BDNF-related expression/signaling. | Literature-constrained direction; calibrated threshold/rates | Smith et al., 1995; Duman & Monteggia, 2006; Tsankova et al., 2006 |
| Eq. 6: Nrf2 | Glucocorticoid-receptor signaling can repress Nrf2-mediated antioxidant responses. | Literature-constrained direction; calibrated threshold/rates | Alam et al., 2017 |
| Eq. 7: Inflammation | Chronic stress and glucocorticoid resistance can support inflammatory activation. | Phenomenological; literature-constrained direction | Dantzer et al., 2008; Miller & Raison, 2016 |
| Eqs. 8-9: Trp/Kyn | Inflammation activates IDO-dependent diversion of tryptophan toward kynurenine. | Literature-constrained structure; calibrated rates | Dantzer et al., 2008; Reus et al., 2015; Miller & Raison, 2016 |
| Eq. 10: Slow stress load | Allostatic concepts motivate cumulative stress exposure; the BDNF multiplier is explicitly hypothesized. | Model-defined integrator with hypothesized feedback gain | McEwen & Stellar, 1993; McEwen, 1998; Numakawa et al., 2009; Suri & Vaidya, 2013 |

*These references motivate the direction and biological plausibility of the couplings. They do not uniquely establish the implemented functional forms or numerical parameter values.*
